# Transcriptional Profiling of D1-MSNs Reveals SIRT1-Dependent and -Independent Responses to Chronic Social Defeat Stress

**DOI:** 10.1101/2025.11.03.686384

**Authors:** Hee-Dae Kim, Hyung Joon Cho, Angel Nguyen, Alyssa Barney, Shenfeng Qiu, Deveroux Ferguson

**Affiliations:** Department of Basic Medical Sciences, University of Arizona College of Medicine-Phoenix, Phoenix, AZ, USA

**Author notes:** Correspondence, Department of Basic Medical Sciences, University of Arizona College of Medicine-Phoenix, Phoenix, AZ, USA.

**Keywords:** SIRT1, Depression, Anxiety, cell-type specificity, translatome, nucleus accumbens

## Abstract

SIRT1 is a critical regulator of neuronal functions and has been implicated in various neurological disorders, including depression. In a previous study, we demonstrated the pro-depressive roles of SIRT1 in the nucleus accumbens (NAc) using the chronic social defeat stress (CSDS) model, a well-validated preclinical mouse model of depression. We found that SIRT1 modulates synaptic functions and exhibits pro-susceptible effects specifically in D1-medium spiny neurons in the NAc. In this study, we extended our investigation of SIRT1 functions by employing cell-type-specific transcriptomics to compare control and susceptible groups under CSDS. Our results revealed that in wild-type mice with functional SIRT1, cellular metabolism is crucial for inducing susceptibility to depression. Conversely, in the absence of functional SIRT1 expression, circadian clock-controlled transcriptional regulation becomes a key factor. These findings suggest that SIRT1 differentially modulates key pathophysiological mechanisms of depression via its deacetylase functions, highlighting its role in both metabolic and circadian regulation in the brain. This research provides new insights into the molecular underpinnings of depression and identifies potential targets for therapeutic intervention.

## Introduction

Depression is a pervasive and debilitating mental disorder affecting millions, with about 20% lifetime prevalence (Kessler et al. 2007). Traditional treatments often fall short, leaving a significant portion of patients with major depressive disorder (MDD) unresponsive (Rush et al. 2006; Trivedi et al. 2006; Fava and Davidson 1996). The complex interplay of genetics and environmental stressors, like chronic stress, is known to trigger MDD (Krishnan and Nestler 2008). The CONVERGE Consortium’s identification of SIRT1, a class III NAD^+^- dependent histone deacetylase (HDAC), as a consistent genetic link to MDD (consortium 2015), supported by various preclinical studies (Abe-Higuchi et al. 2016; Kim et al. 2016; Kim et al. 2024; Libert et al. 2011), marks a significant advance, considering SIRT1’s role in neurodevelopment, aging, addiction, and mental health (Herskovits and Guarente 2014; Lu et al. 2018; Ferguson et al. 2013; Koo et al. 2016; Renthal et al. 2009; Kim et al. 2016; Kim et al. 2024).

This study focuses on the nucleus accumbens (NAc), a key brain reward area comprising two types of medium spiny neurons (D1-MSNs or D2-MSNs), and examines how stress and SIRT1 knockout influence transcriptional changes in D1-MSNs. Using a triple transgenic mouse model to knock out SIRT1 specifically in D1-MSNs and employing the RiboTag approach for precise mRNA isolation (Sanz et al. 2009), we dissected the molecular contributions of stress and SIRT1 deletion on gene expression in D1-MSNs. This targeted approach allows us to unravel the interplay between SIRT1 and stress-induced transcriptional alterations, providing insights into the underlying mechanisms of depression-like behaviors. Our study design involves comparing gene expression profiles of wild-type (WT) and SIRT1 knockout (KO) mice under control (Con) and susceptible (Sus) conditions following chronic social defeat stress (CSDS) (Krishnan et al. 2007). We performed a comprehensive bioinformatic analysis to identify differentially expressed genes and pathways between multiple conditions.

Gene enrichment analysis revealed a significant downregulation of genes involved in mitochondrial function and cellular energy metabolism in D1-MSNs of WT Sus mice. Conversely, the deletion of SIRT1 in D1-MSNs significantly altered circadian clock-regulated genes, which are known to be disrupted in depression (Li et al. 2013; McCarthy et al. 2019; Landgraf et al. 2016). Here, we propose that SIRT1 acts as a critical regulator of cellular metabolism and circadian rhythm-related gene expressions in D1-MSNs, and its deletion may lead to transcriptional alterations that underlie the development of depression-like behaviors.

## Materials and Methods

### Mice

Male C57BL/6J mice (7–9 weeks old) were obtained from Jackson Laboratory and housed on a 12 hr light-dark cycle with access to food and water ad libitum. Male CD1 retired breeder mice (9–13 months old) were obtained from Charles River Laboratories. Mice acclimated to the facility for 1 week before any experimentation. To induce deletion of the functional Sirt1 transcript in D1 neurons in the NAc, homozygote Sirt1 floxed mice (RRID:IMSR_JAX:008041) (Li et al. 2007) crossed with D1-Cre hemizygote (line FK150) BAC transgenic mice (mice expressing Cre-recombinase driven by the dopamine receptor 1 promoter) from GENSAT (Gong et al. 2007) on a C57BL/6J background were used for behavioral experiments. Results from this cross generated mice lacking a functional copy of the SIRT1 gene specifically in D1-MSNs, hereafter referred to as SIRT1^D1-KO^ mice displayed no gross anatomical or reproductive defects and were born with normal mendelian ratios. To induce Rpl22-HA in D1 neurons, we crossed homozygous RiboTag mice (RRID:IMSR_JAX:029977) (Sanz et al. 2009) with D1 hemizygote mice. All behavioral tests and animal sacrifices were performed at zeitgeber time (ZT) 02-06. All animal procedures were approved by the University of Arizona Medical School Institutional Animal Care and Use Committees.

### Chronic social defeat stress

Social defeat stress was performed according to published protocols (Golden et al. 2011; Kim et al. 2017). Test mice were exposed to an unfamiliar and aggressive (less than 1-minute latency to start aggressive behaviors) male CD1 retired breeder mouse for 5 min per day for up to 10 days. Following direct interaction with the CD1 aggressor, animals were placed in an adjacent compartment of the same cage for the next 24 hrs with sensory, but not physical contact. Control animals were housed in equivalent cages but with members of the same strain. 24 hrs following the last social defeat, animals were assayed on the social interaction test and sorted into either susceptible or unsusceptible (resilient) phenotypes based on interaction scores. Briefly, for the social interaction test in the first 2.5 min trial (“target absent”), the test mouse was allowed to freely explore a square-shaped open-field arena (44 x 44 cm) possessing a removable perforated plexiglass enclosure (10 x 6 cm) opposed to one side. During the second 2.5 min trial (“target present”), the mouse was reintroduced into this arena now containing an unfamiliar CD1 mouse within the smaller cage. EthoVision video-tracking software (Noldus, Leesburg, VA) was employed to measure time spent in the “interaction zone” (14 x 26 cm). The social interaction (SI) ratio was calculated as 100 x (interaction zone time of “target present”)/(interaction zone time of “target absent”) (Golden et al. 2011; Kim et al. 2017; Krishnan et al. 2007). Mice showing less than 100% of SI ratio are grouped as depression-susceptible mice, and mice showing over 100% SI ratio belong to depression-resilient group.

### Immunohistochemistry

Mice were given a lethal dose of Euthasol (Virbac, Westlake, TX), then sequentially perfused with PBS and 4% paraformaldehyde. Brains were post-fixed in 4% paraformaldehyde overnight and cryoprotected in 30% sucrose solution at 4°C. After equilibration in the sucrose solution, brains were sectioned at 30 μm on a sliding block microtome (American Optical). Free-floating sections were washed with PBS, then blocked (3% donkey serum, 0.1% Triton X-100 in PBS). The following primary antibodies were applied overnight at 4°C with constant agitation: anti-CRE (mouse monoclonal, 1:500; Millipore, Burlington, MA), and anti-HA (rabbit polyclonal, 1:2000; Abcam). After 4°C overnight incubation with constant agitation, sections were incubated with appropriate secondary antibodies conjugated with fluorescent dyes (Alexa 488 and Alexa 555; Life technologies) for 2 hr at room temperature. They were subsequently washed and mounted with Vectashield containing DAPI (Vector Labs, Burlingame, CA) and analyzed using confocal microscopy employing a 20x objective (LSM710; Carl Zeiss, San Diego, CA).

### RiboTag immunoprecipitation

Six NAc tissue punches (2 mm) from each SIRT1^D1-KO^/Rpl22 or D1-Cre/Rpl22 mouse are homogenized in 1 ml of homogenization buffer for each sample (2 mice, which have similar SI ratio, pooled for each sample). Samples are then centrifuged at 4°C, 10,000 g for 10 min and 800 μl of clear supernatants are mixed with 5 μg of HA- antibody (Abcam, Cambridge, MA), and incubated on a gentle rotator at 4°C overnight. The next day, 200 μl of magnetic beads (Dynabeads anti-rabbit IgG; Life technology, Carlsbad, CA) are added and incubated overnight with constant rotation. The following day, magnetic beads are washed 3 times in the magnetic rack for 10 min with a high salt wash buffer. Each RNA sample was extracted using Direct-zol miniprep kit (Zymo Research, Irvine, CA) and subjected to qRT-PCR or RNA-seq. For quality control, RNA integrity (RIN) values were measured and samples with RIN < 7 were excluded.

### Library preparation and RNA-seq

RNA-seq was performed by UAGC (University of Arizona Genetics Core, University of Arizona, Tucson, AZ). Briefly, purified RNA samples from RiboTag immunoprecipitation (500 ng of RNA for each) were prepped for libraries after poly-A selection (KAPA Stranded mRNA-Seq Kit; Roche, Indianapolis, IN) and the libraries were sequenced in NextSeq500 (Illumina, San Diego, CA). Randomly chosen samples were divided into four sets and run in high output mode (maximum 400 million reads, 2 x 150bp paired end reads) for each set. All the samples were normalized before pooling and loading on the sequencer.

### RNA-seq and statistical data analysis

Data was analyzed by Rosalind (https://rosalind.onramp.bio/), with a HyperScale architecture developed by OnRamp BioInformatics, Inc. (San Diego, CA). Reads were trimmed using cutadapt. Quality scores were assessed using FastQC. Reads were aligned to the Mus musculus genome (mm10) using STAR. Individual sample reads were quantified using HTseq and normalized via Relative Log Expression (RLE) using DESeq2 R library. Read distribution percentages, violin plots, identity heatmaps, and sample multidimensional scaling (MDS) plots were generated as part of the QC step using RSeQC. DEseq2 was also used to calculate fold changes and p-values. Clustering of genes for the final heatmap of differentially expressed genes (DEGs) was done using the Partitioning Around Medoids (PAM) method using the fpc R library. Functional enrichment analysis of pathways, gene ontology, domain structure, and other ontologies was performed using HOMER. Several database sources were referenced for enrichment analysis, including Interpro, NCBI, KEGG, MSigDB, REACTOME, and WikiPathways. Enrichment was calculated relative to a set of background genes relevant to the experiment. Additional gene enrichment analyses were performed using Enrichr (Kuleshov et al. 2016). GO and KEGG pathway plots were generated with SRplot (Tang et al. 2023). Venn diagrams were created with a web-based tool (http://bioinformatics.psb.ugent.be/webtools/Venn/). Additional statistical analyses were conducted using Prism (version 9.5.0; GraphPad, San Diego, CA).

## Results

In our previous study, we reported that D1- and D2-MSNs show distinct gene expression profiles in chronic stress conditions (Kim et al. 2021). Here we analyzed both WT and D1-MSN specific Sirt1-KO (SIRT1^D1-KO^) mice that underwent CSDS and compared the expression profiles of control (Con) and susceptible (Sus) groups in the WT and KO mice. To do that, we employed cell-type specific transcriptomics using RiboTag crossed with D1-Cre or SIRT1^D1-KO^ mice while performing CSDS (Fig. 1a). Based on the RiboTag immunoprecipitation (IP) of D1-specific transcripts (Fig. 2b), RNA sequencing shows enrichment of coding regions, indicating mature mRNAs, in isolated transcripts compared to input samples (Fig. 1c; Table S1). The transcriptome profiles exhibit distinct clustering between WT and KO samples, which was not observed in the input samples (Fig. 1d). We further validated the D1-specific profiles with marker genes in both WT and KO samples (Fig. 1e).

**Fig. 1.**
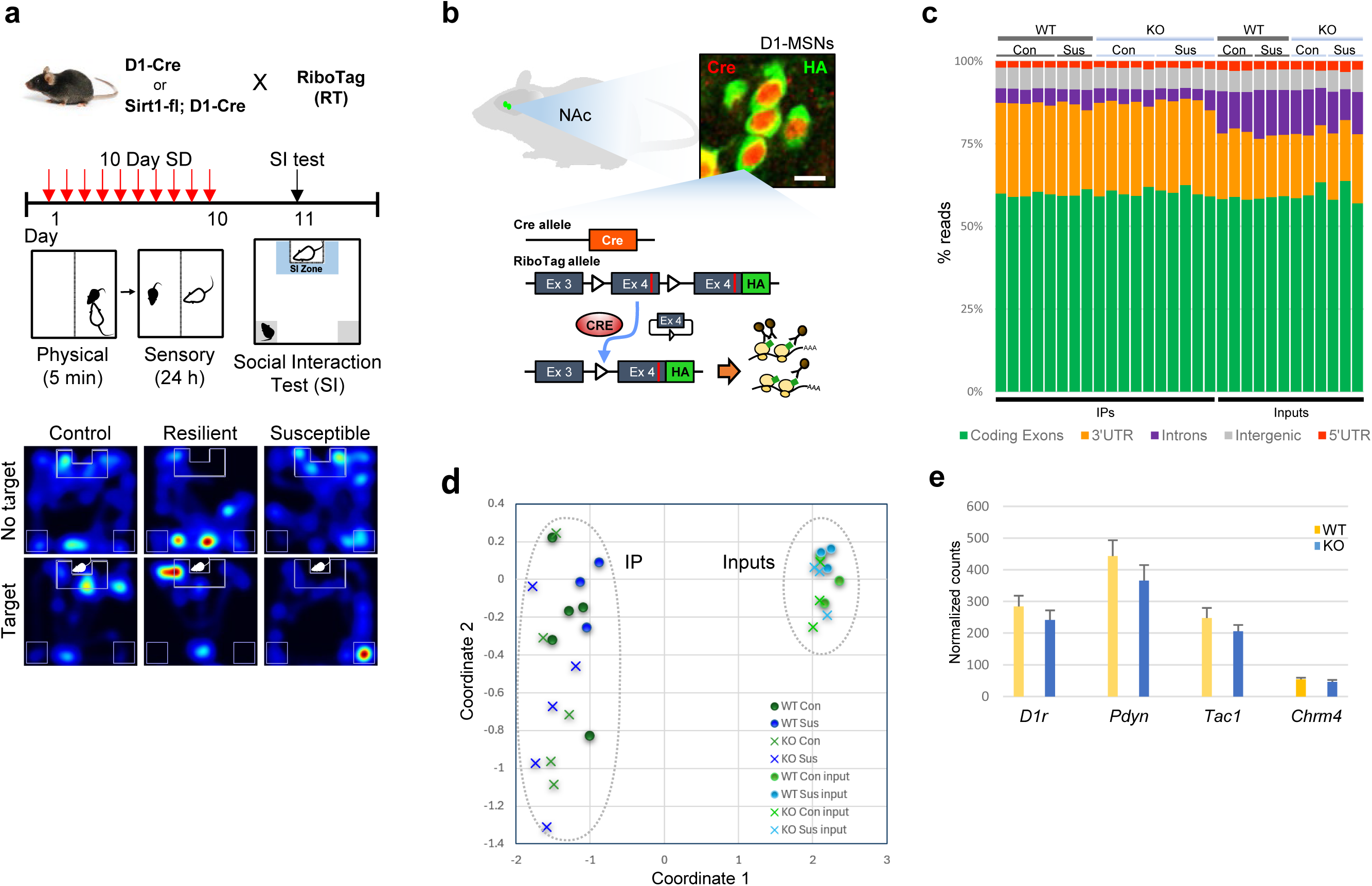
Experimental model and isolation of D1-MSN-specific transcriptomics using RiboTag. **a** Generation of D1-MSN-specific Sirt1-knockout model (SIRT1^D1-KO^) crossed with RiboTag. The mice underwent chronic social defeat stress. The representative heatmap shows the results of the social interaction (SI) test for control, resilient, and susceptible mice. **b** Schematics of RiboTag-based cell-type specific transcriptomics and immunohistochemical validation of the RiboTag model (NAc, coronal; scale bar, 10 μm). **c** Read mapping composition of each RNA-seq sample on the mouse genome, mm10 (% average for coding regions: IPs, 60.0% vs inputs, 59.3%; for intronic regions: IPs, 4.4% vs inputs, 12.6%). **d** Clustering of RNA-seq samples of wild-type and Sirt1-knockout mice in a multidimensional scaling plot. **e** Normalized expression levels of D1-MSN specific marker genes, D1r, Pdyn, Tac1, and Chrm4

**Fig. 2.**
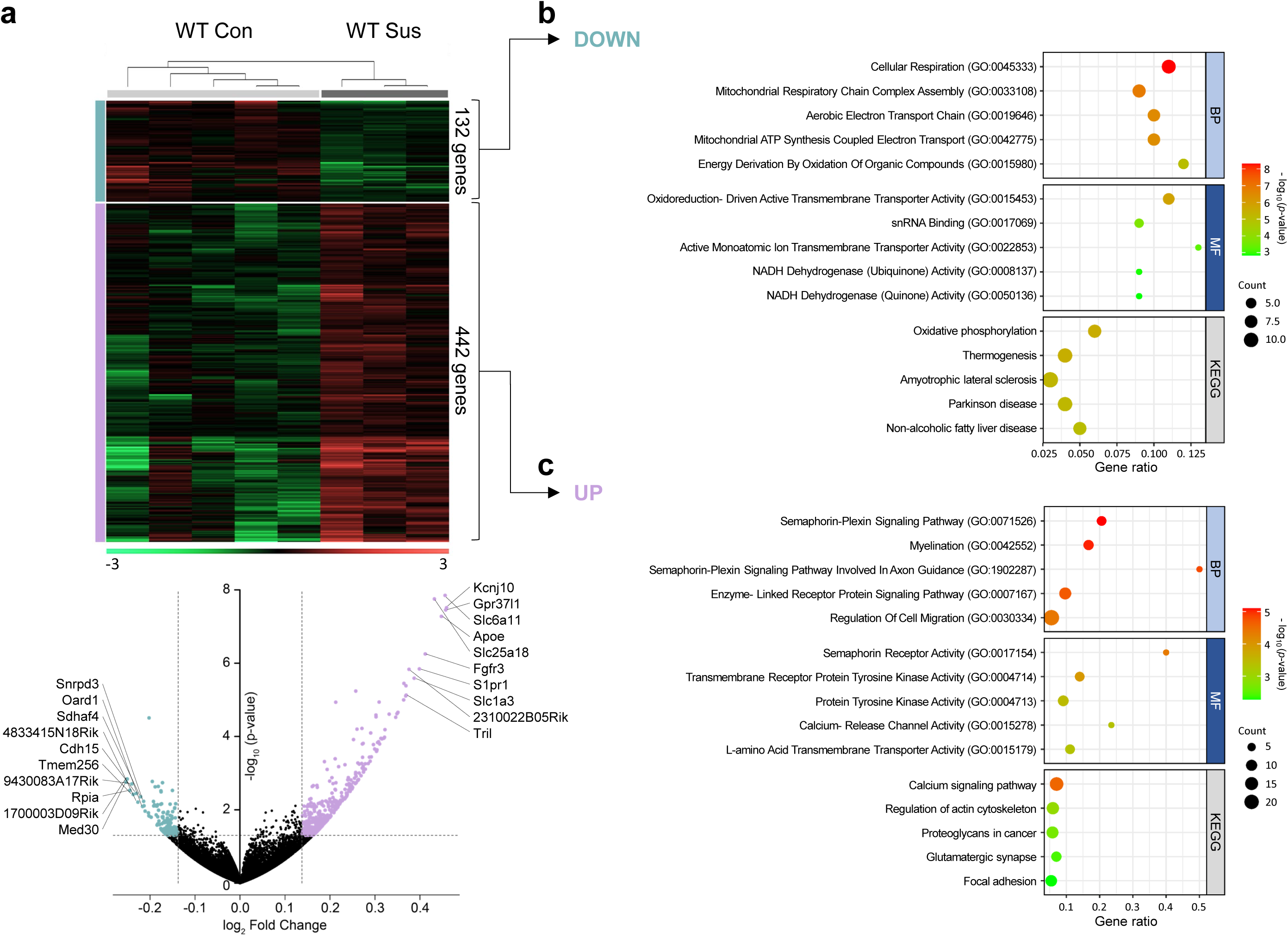
Analysis of susceptible mice-specific genes in wild-type D1-MSNs. **a** The gene expression differences between wild-type control (WT Con) and susceptible (WT Sus) mice are presented as a heatmap and volcano plot, showing 574 differentially expressed genes. In the comparison between WT Con and WT Sus, 132 genes were decreased and 442 genes were increased in WT Sus, using thresholds of a fold change of ±1.1 and a *p*-value < 0.05. **b, c** Gene ontology (GO) analysis of the downregulated (**b**) and upregulated (**c**) DEGs was performed using EnrichR, focusing on biological process (BP), KEGG pathway, and molecular function (MF)

To determine the baseline effect of stress on gene expression in D1-MSNs when the SIRT1 pathway is intact, we performed differential gene expression analysis between WT Con (n=5) and WT Sus (n=3) immunoprecipitated transcripts, a heatmap and volcano plot were generated to visualize the patterns and significance of gene expression differences. A total of 574 differentially expressed genes (DEGs) were identified, with 442 genes found to be upregulated and 132 downregulated (*p*<0.05) in the WT Sus group (Fig. 2a; Table S2). The heatmap (Fig. 2a, top) displays the clustering of expression patterns, showing distinct expression profiles between WT Con and WT Sus, with the degree of expression indicated by the color gradient from green (low expression) to red (high expression). In the accompanying volcano plot (Fig. 2a, bottom), the DEGs plotted with fold change on the x-axis and −log10 *p*-value on the y-axis, highlighting upregulated genes in WT Sus relative to WT Con in purple and downregulated genes in green. The top most significantly upregulated genes in the WT Sus group, ranked by fold change and *p*-value, were Kcnj10 (1.37-fold, *p* = 3.07e-08), Gpr37l1 (1.37-fold, *p* = 3.53e-08), Slc6a11 (1.37-fold, *p* = 1.42e-08), Apoe (1.36-fold, *p* = 5.29e-08), Slc25a18 (1.35-fold, *p* = 1.77e-08), Fgfr3 (1.33-fold, *p* = 5.60e-07), S1pr1 (1.32-fold, *p* = 1.44e-06), Slc1a3 (1.31-fold, *p* = 2.58e-06), 2310022B05Rik (1.30-fold, *p* = 1.48e-06), and Tril (1.29-fold, *p* = 7.49e-06). These genes are involved in various biological processes, including ion transport (Kcnj10, Slc6a11, Slc25a18, Slc1a3), G protein-coupled receptor signaling (Gpr37l1, S1pr1), lipid metabolism (Apoe), growth factor signaling (Fgfr3), and immune responses (Tril). Conversely, the genes that demonstrated the most significant decrease in expression in WT Sus compared to WT Con are Med30 (−1.19-fold, *p* = 1.76e-03), 1700003D09Rik (−1.19-fold, *p* = 1.44e-03), Rpia (−1.19-fold, *p* = 2.96e-03), 9430083A17Rik (−1.18-fold, *p* = 1.97e-03), Tmem256 (−1.18-fold, *p* = 3.81e-03), Cdh15 (−1.17-fold, *p* = 3.57e-03), 4833415N18Rik (−1.17-fold, *p* = 6.17e-03), Sdhaf4 (−1.17-fold, *p* = 4.40e-03), Oard1 (−1.16-fold, *p* = 8.11e-03), and Snrpd3 (−1.16-fold, *p* = 5.87e-03). These genes are associated with various cellular functions, such as transcriptional regulation (Med30), energy and metabolic functions (Rpia, Sdhaf4, Oard1), cell adhesion (Cdh15), and RNA splicing (Snrpd3).

Gene enrichment analysis using EnrichR (Kuleshov et al. 2016) revealed significant downregulation of genes involved in several biological processes associated with mitochondrial function and cellular energy metabolism in WT Sus D1-MSNs. Specifically in the gene ontology (GO) for Biological Process (Fig. 2b), we observed a downregulation in the expression of genes involved in ‘Cellular Respiration’ (GO:0045333, −log_10_(*p*) = 8.31) which suggests a compromised basic energy-producing capacity in WT Sus mice. The ‘Mitochondrial Respiratory Chain Complex Assembly’ (GO:0033108, −log_10_(*p*) = 6.84) and ‘Mitochondrial ATP Synthesis Coupled Electron Transport’ (GO:0042775, −log_10_(*p*) = 6.40) gene sets were also significantly reduced. These findings suggest a possible attenuation of ATP production efficiency in the mitochondria of D1-MSNs from WT Sus mice. Gene sets upregulated in D1-MSNs WT Sus mice under Biological Process GO, revealed a particular emphasis on the dysregulation of pathways involved in neuronal development and signaling, the ‘Semaphorin-Plexin Signaling Pathway’ (GO:0071526), with the most significant −log_10_(*p*) = 5.10 (Fig. 2c). Its significant upregulation suggests a restructuring of neuronal circuits in response to chronic stress (Tran, Kolodkin, and Bharadwaj 2007). ‘Myelination’ (GO:0042552) was the next enriched category, with a −log_10_(*p*) = 5.03. Myelination is critical for the proper functioning of the nervous system, affecting signal transmission speed and neural plasticity. Stress-induced myelination changes may contribute to cognitive and emotional deficits in stress-related disorders (Fields 2008).

The key Molecular Functions GO that was decreased, predominantly affected transmembrane transport activity and redox processes. The most significantly downregulated function was ‘Oxidoreduction-Driven Active Transmembrane Transporter Activity’ (GO:0015453, −log_10_(*p*) = 5.83), suggesting a reduced capability of transmembrane proteins to couple redox reactions with the active transport of molecules across cellular membranes (Fig. 2b). Moreover, the gene sets associated with ‘NADH Dehydrogenase (Ubiquinone) Activity’ (GO:0008137, −log_10_(*p*) = 2.87) and ‘NADH Dehydrogenase (Quinone) Activity’ (GO:0050136, −log_10_(*p*) = 2.83) were considerably downregulated (Fig. 2b). NADH dehydrogenase is the largest complex within the mitochondrial respiratory chain, and its reduced activity implies a potential compromise in mitochondrial function and ATP synthesis, which are critical for energy-demanding processes in neurons (Breuer et al. 2013). The Molecular Function GO that were increased following stress was ‘Semaphorin Receptor Activity’ (GO:0017154; −log_10_(*p*) = 4.35) and ‘Transmembrane Receptor Protein Tyrosine Kinase Activity’ (GO:0004714; −log_10_(*p*) = 3.46) (Fig. 2c). KEGG analysis identified significant downregulation of gene sets associated with ‘Oxidative Phosphorylation’ (−log_10_(*p*) = 5.54) and ‘Thermogenesis’ (−log_10_(*p*) = 5.48) in WT Sus mice after CSDS as compared to WT Con mice (Fig. 2b). The downregulation of ‘Oxidative Phosphorylation’ is particularly compelling, as this pathway is critical for energy production. A decrease in genes related to this pathway may reflect a lowered metabolic efficiency and could be indicative of mitochondrial dysfunction. In genes that were increased, there were enrichments in the ‘Calcium Signaling Pathway’ (−log_10_(*p*) = 4.57), the ‘Regulation of Actin Cytoskeleton’ (−log_10_(*p*) = 2.92), and the ‘Glutamatergic Synapse’ (−log_10_(*p*) = 2.41) pathways in WT Sus mice (Fig. 2c). The significant enrichment of these pathways suggests that chronic stress might lead to heightened intracellular calcium levels, which could contribute to altered neuronal function (Berridge 2012).

To understand how stress affects gene expression in D1-MSNs without SIRT1, we performed DEG analysis on two groups: SIRT1^D1-KO^ control (KO Con, n=5) and SIRT1^D1-KO^ susceptible mice (KO Sus, n=5). Analyzing these groups identifies stress-regulated genes independent of SIRT1 in D1-MSNs, which may reveal alternative mechanisms by which stress alters gene regulation in these neurons. A distinct expression profile was observed, with 90 DEGs identified. Of these, 39 were upregulated and 51 downregulated in the KO Sus group compared to KO Con (Fig. 3a; Table S3). The top upregulated genes in KO Sus relative to KO Con, ranked by fold change, were Aldh7a1 (1.20-fold, *p* = 1.42e-05), Dnajc21 (1.15-fold, *p* = 6.07e-04), Bhlhe40 (1.15-fold, *p* = 1.53e-03), D030028A08Rik (1.14-fold, *p* = 3.74e-03), Pcdhgb6 (1.14-fold, *p* = 3.91e-03), AI846148 (1.13-fold, *p* = 2.77e-03), Ubtd1 (1.12-fold, *p* = 9.23e-03), Caap1 (1.12-fold, *p* = 9.83e-03), P4ha1 (1.12-fold, *p* = 5.02e-03), and Etv6 (1.12-fold, *p* = 8.80e-03). These genes are involved in various biological processes, including metabolism (Aldh7a1, P4ha1), RNA processing or ribosome biogenesis (Dnajc21), regulation of cellular processes including circadian rhythm (Bhlhe40, Etv6), cell-cell connections in the brain (Pcdhgb6), and cell cycle regulation (Caap1). The top downregulated genes in KO Sus included Parvb (−1.16-fold, *p* = 1.10e-03), Snhg11 (−1.16-fold, *p* = 3.49e-04), Mzf1 (−1.15-fold, *p* = 1.43e-03), 2700049A03Rik (−1.15-fold, *p* = 1.96e-03), 2610005L07Rik (−1.14-fold, *p* = 3.28e-03), Gas2 (−1.14-fold, *p* = 3.88e-03), Hdac7 (−1.14-fold, *p* = 4.15e-03), L3mbtl1 (−1.13-fold, *p* = 3.77e-03), Reck (−1.13-fold, *p* = 4.09e-03), and Pdgfrb (−1.13-fold, *p* = 2.29e-03). These genes are implicated in various functions, including regulation cytoskeleton and cell structure (Parvb, Gas2), cell migration, proliferation, and adult neurogenesis (Snhg11) chromatin organization and transcriptional regulation (Mzf1, Hdac7, L3mbtl1), and cell differentiation and angiogenesis (Reck, Pdgfrb).

**Fig. 3.**
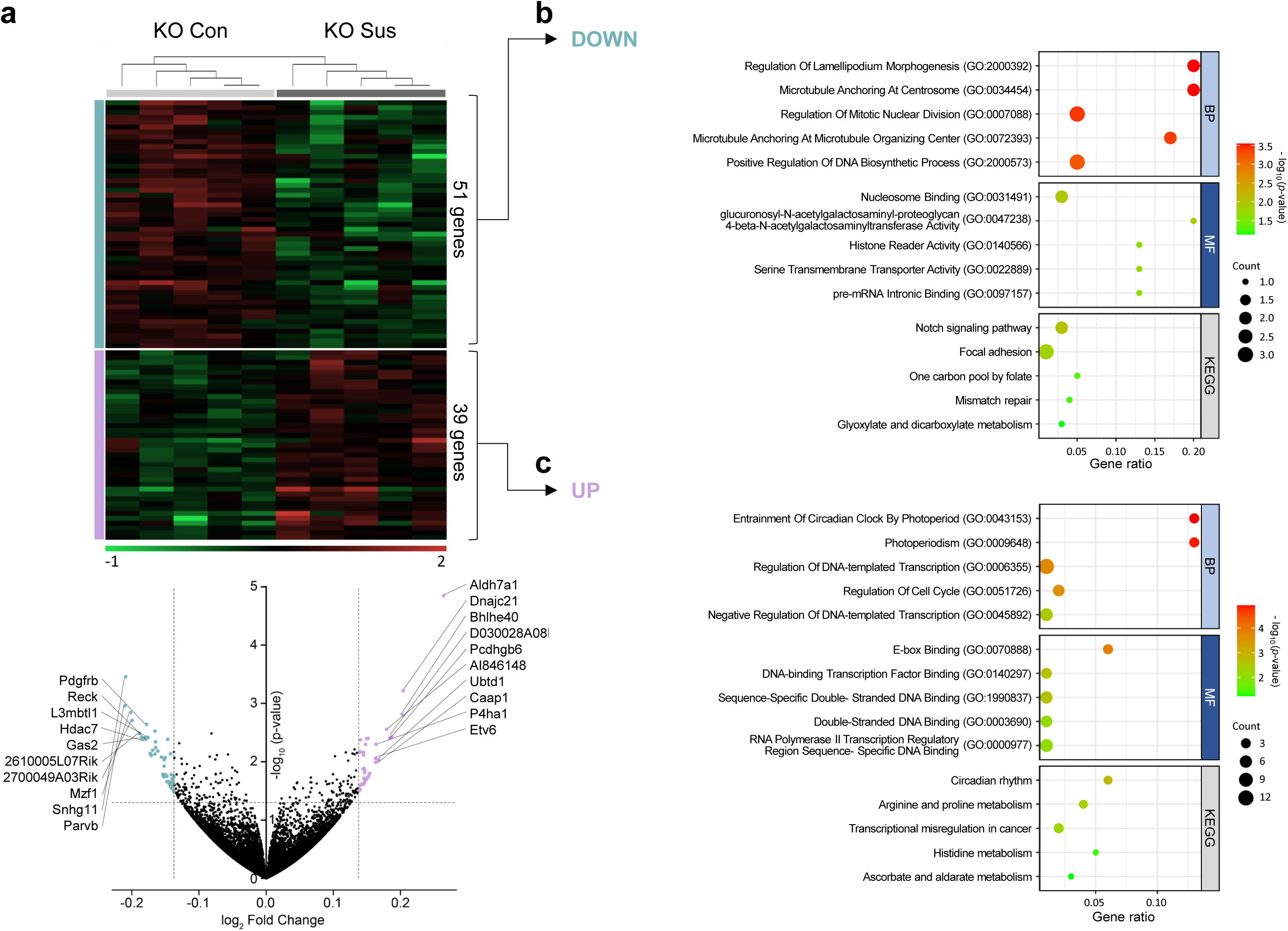
Analysis of susceptible mice-specific genes in SIRT1-KO D1-MSNs. **a** The gene expression differences between SIRT1-KO control (KO Con) and susceptible (KO Sus) mice are presented as a heatmap and volcano plot, showing 90 differentially expressed genes. In the comparison between KO Con and KO Sus, 51 genes were decreased and 39 genes were increased in KO Sus, using thresholds of a fold change of ±1.1 and a *p*-value < 0.05. **b, c** Gene ontology (GO) analysis of the downregulated (**b**) and upregulated (**c**) genes was performed using EnrichR, focusing on biological process (BP), KEGG pathway, and molecular function (MF)

Overall GO analysis of downregulated and upregulated genes in SIRT1^D1-KO^ susceptible mice (Fig. 3b, c) revealed significant enrichment in processes related to circadian rhythm regulation. The most significant Biological Process GO term, ‘Entrainment of Circadian Clock by Photoperiod’ (GO:0043153, −log_10_(*p*) = 4.92), suggests an enhanced synchronization of the circadian clock with natural light-dark cycles in these animals (Fig. 3c). Additionally, the term ‘Photoperiodism’ (GO:0009648, −log_10_(*p*)= 4.86) was highly enriched, indicating a heightened biological response to the length of daylight, which is crucial for orchestrating seasonal behaviors. These findings highlight the potential role of SIRT1 in modulating the circadian clock and photoperiodic responses in the context of stress susceptibility. In support of the significant effect of SIRT1 deletion on the expression of circadian-related genes, the Molecular Function GO revealed ‘E-Box Binding’ (GO:0070888, −log_10_(*p*)= 3.87) and ‘DNA-binding Transcription Factor Binding’ (GO:0140297, −log_10_(*p*)= 2.66) as top regulated functions (Fig. 3c). E-box Binding refers to the interaction with E-box motifs, which are specific DNA sequences (CANNTG) recognized by transcription factors, notably within the basic helix-loop-helix family. These motifs are often found in the promoter regions of genes that regulate circadian rhythm and development. Given that SIRT1 interacts with the circadian molecular circuitry by deacetylating key transcription factors like CLOCK, an increase in E-box binding activity suggests a possible compensatory response to maintain circadian gene expression in the absence of SIRT1’s regulation (Hirayama et al. 2007). The heightened binding at E-box motifs in susceptible animals may be indicative of altered transcriptional activity of genes that are critical for the adaptation to environmental stressors. KEGG pathways analysis confirmed these results, with the most enriched term ‘Circadian rhythm’ (−log_10_(*p*)=2.77), suggesting stress and SIRT1 deletion may alter circadian clock gene expression in D1-MSNs (Fig. 3c). An analysis of gene ontology within the Molecular Function domain reveals a significant decrease in ‘Nucleosome Binding’ (GO:0031491, −log_10_(*p*) = 1.96) and ‘Histone Reader Activity’ (GO:0140566, −log_10_(*p*)= 1.69), which suggests alterations in chromatin structure and dynamics, which are vital for the regulation of gene expression (Fig. 3b) (Jenuwein and Allis 2001). The observed decreases in these molecular functions in the context of SIRT1 deletion align with previous reports highlighting SIRT1’s involvement in chromatin remodeling, gene expression regulation, and neural plasticity (Hisahara et al. 2008; Michan et al. 2010).

To assess the baseline effect of SIRT1 KO on gene expression in D1-MSNs, we performed differential gene expression analysis between wild-type control (WT CON, n=5) and SIRT1^D1-KO^ control (KO CON, n=5). The analysis identified total 792 DEGs, with 339 genes upregulated and 454 genes downregulated (Fig. 4a; Table S4). The top genes with the most pronounced increase in expression in KO CON, ranked by fold change, were Gm1943 (2.04-fold, *p* = 2.05e-27), A930005H10Rik (1.52-fold, *p* = 1.57e-10), Cox18 (1.51-fold, *p* = 1.40e-11), Adi1 (1.50-fold, *p* = 5.78e-10), Cfap54 (1.48-fold, *p* = 1.62e-09), 1500015A07Rik (1.43-fold, *p* = 2.74e-08), Stbd1 (1.41-fold, *p* = 3.51e-10), A930033H14Rik (1.41-fold, *p* = 7.43e-08), Gm14305 (1.40-fold, *p* = 8.35e-08), and Hddc3 (1.38-fold, *p* = 1.01e-06). These genes participate in key biological processes such as mitochondrial function and metabolism (Cox18, Adi1, Stbd1) and the operation of cilia and flagella (Cfap54). However, many of the top-upregulated genes remain less characterized.

**Fig. 4.**
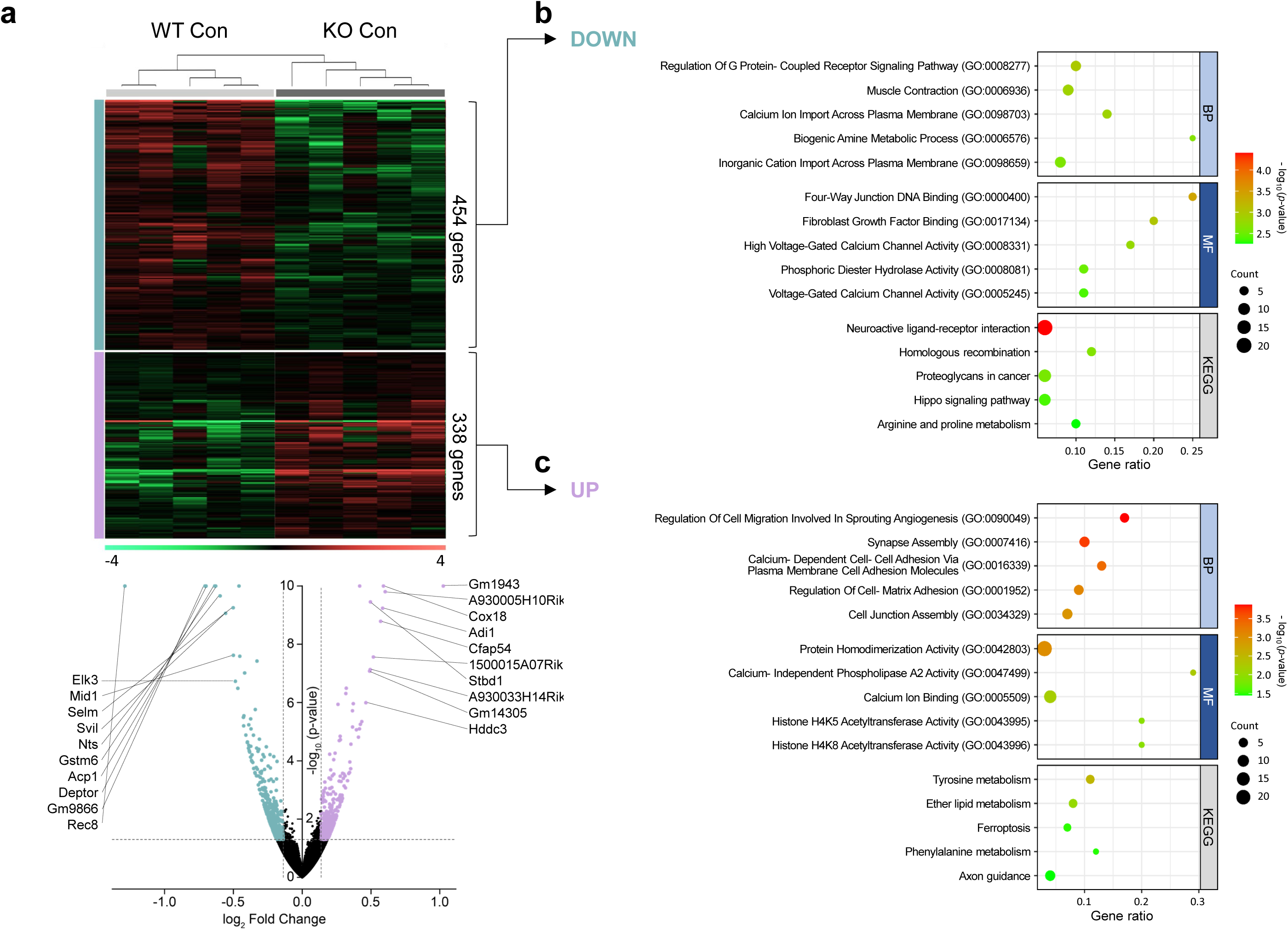
Analysis of SIRT1-KO mice-specific genes in D1-MSNs of control mice. **a** The gene expression differences between wild-type control (WT Con) and SIRT1-KO control (KO Con) mice are presented as a heatmap and volcano plot, showing 792 differentially expressed genes. In the comparison between WT Con and KO Con, 454 genes were decreased and 338 genes were increased in KO Con, using thresholds of a fold change of ±1.1 and a *p*-value < 0.05. **b, c** Gene ontology analysis of the downregulated (**b**) and upregulated (**c**) genes was performed using EnrichR, focusing on biological process (BP), KEGG pathway, and molecular function (MF)

The genes exhibiting the most significant decrease in expression included Rec8 (−2.44-fold, *p* ≈ 0), Gm9866 (−1.63-fold, *p* = 3.88e-19), Deptor (−1.62-fold, *p* = 3.42e-18), Acp1 (−1.55-fold, *p* = 6.84e-15), Gstm6 (−1.55-fold, *p* = 2.71e-11), Nts (−1.51-fold, *p* = 2.18e-10), Svil (−1.47-fold, *p* = 8.65e-10), Selm (−1.42-fold, *p* = 5.57e-10), Mid1 (−1.41-fold, *p* = 2.40e-08), and Elk3 (−1.40-fold, *p* = 1.87e-07). These genes are involved in several biological functions, such as chromatin remodeling and gene regulation (Rec8, Mid1, Elk3), metabolic functions (Acp1, Gstm6, Selm), signal transduction and neurotransmission (Deptor, Nts), and cytoskeletal organization (Svil).

GO analysis for the Biological Processes revealed that SIRT1 deletion in D1-MSNs significantly downregulated genes in the ‘Regulation of G Protein-Coupled Receptor Signaling Pathway’ (GO:0008277) with (−log_10_(*p*) = 2.97) (Fig. 4b). This suggests that the absence of SIRT1 substantially impacts GPCR-mediated signaling events in these neurons. Another notable result was the decrease in ‘Calcium Ion Import Across Plasma Membrane’ (GO:0098703) with a (−log_10_(*p*)=2.84) (Fig. 4b). Calcium ions play a crucial role in various cellular processes, including neurotransmission, and a reduction in their transport into cells could have multiple downstream consequences. From the Molecular Function GO of genes downregulated the most significantly enriched term, ‘Four-Way Junction DNA Binding’ (GO:0000400), suggests that SIRT1 deletion leads to reduced expression of genes involved in binding and resolving Holliday junctions, which are crucial intermediates in DNA recombination and repair (Fig. 4b) (Schwacha and Kleckner 1995). This aligns with SIRT1’s known role in maintaining genomic stability and regulating DNA repair processes (Oberdoerffer et al. 2008). Interestingly, two of the top five terms relate to calcium channel activity: ‘High Voltage-Gated Calcium Channel Activity’ (GO:0008331) and ‘Voltage-Gated Calcium Channel Activity’ (GO:0005245) (Fig. 4b). This indicates that the loss of SIRT1 in D1-MSNs under basal conditions results in downregulation of genes encoding calcium channels, particularly those activated by high voltages. Voltage-gated calcium channels are critical for regulating neuronal excitability, synaptic transmission, and plasticity in the brain (Catterall 2011). KEGG analysis revealed that ‘Neuroactive ligand-receptor interaction’ was the most significantly enriched pathway (Fig. 4b), which plays a pivotal role in synaptic transmission and neural communication. The downregulation of this pathway in the KO Con animals suggests that SIRT1 profoundly affects the capacity of neurons to respond to neurotransmitters and neuromodulators. ‘Homologous recombination’ term was also significantly enriched −log_10_(p)= 2.64 and is essential for the repair of DNA double-strand breaks and the maintenance of genomic stability (Fig. 4b). The observed downregulation implies that SIRT1 might play a role in preserving genome integrity in neuronal cells, consistent with previous findings that SIRT1 participates in DNA damage response pathways (Oberdoerffer et al. 2008). Reduced efficacy of this pathway could lead to an accumulation of genetic damage over time, which could be particularly detrimental to long-lived cells such as neurons.

Genes that were upregulated after SIRT1 deletion showed enrichment of ‘Synapse Assembly’ (GO:0007416, −log_10_(*p*) = 3.71) and ‘Calcium-Dependent Cell-Cell Adhesion Via Plasma Membrane Cell Adhesion Molecules (CAMs)’ (GO:0016339, −log_10_(*p*) = 3.37) (Fig. 4c). Synapse assembly is a crucial process in neurodevelopment and synaptic plasticity, enabling proper neuronal connectivity and communication (Garner et al. 2002). The increased expression of synapse assembly-related genes in SIRT1^D1-KO^ mice implies that SIRT1 plays a role in regulating synaptic structure and function in D1-MSNs (Kim et al. 2024). In the nervous system, CAMs play essential roles in neurite outgrowth, synapse formation, and synaptic plasticity (Shapiro, Love, and Colman 2007). The upregulation of genes related to calcium-dependent cell-cell adhesion suggests that SIRT1 may be involved in modulating the expression of CAMs, potentially influencing neuronal connectivity and synaptic function. The Molecular Function GO of upregulated genes revealed an interesting finding of enrichment of histone H4K5 and H4K8 acetyltransferase activity (GO:0043995 and GO:0043996) with −log_10_(*p*) = 1.93 for both (Fig. 4c). The increased expression of genes with histone H4K5 and H4K8 acetyltransferase activities in SIRT1^D1-KO^ D1-MSNs suggests that SIRT1 regulates histone acetylation levels and chromatin structure in these neurons. This observation is consistent with the known function of SIRT1 as a histone deacetylase, modulating gene expression through chromatin modifications (Vaquero et al. 2004; Imai et al. 2000). KEGG analysis revealed a notable enrichment in ‘Tyrosine metabolism’ with a −log_10_(*p*) = 2.51, indicating a substantial shift away from normal physiological conditions (Fig. 4c). This pathway is integral to the synthesis of key neurotransmitters and has been implicated in various neurological disorders when dysregulated. Similarly, ‘Axon guidance’ was also upregulated (Fig. 4c), which is crucial for the development and repair of neuronal networks. This suggests potential compensatory neural plasticity or maladaptive changes in response to the absence of SIRT1, which is known to play a role in neuroprotection and plasticity.

To evaluate stress response differences in SIRT1^D1-KO^ mice, we conducted a differential gene expression analysis comparing wild-type susceptible (WT Sus, n=3) and SIRT1^D1-KO^ susceptible (KO Sus, n=5) mice. This analysis identifies how SIRT1 deletion impacts gene expression in response to stress, highlighting differentially regulated pathways in wild-type and Sirt1 knockout mice subjected to CSDS. The analysis identified 2328 DEGs, with 1016 genes upregulated and 1312 genes downregulated (Fig. 5a; Table S5). Clear differentiation in gene expression levels is also evident in the clustering patterns observed in the heatmap. The top upregulated genes in KO Sus, ranked by fold change, were Adi1 (2.19-fold, *p* = 7.40e-26), Hddc3 (1.96-fold, *p* = 5.74e-20), Cfap54 (1.81-fold, *p* = 2.77e-09), Erich5 (1.78-fold, *p* = 5.59e-08), Stbd1 (1.77-fold, *p* = 1.02e-08), Gm1943 (1.68-fold, *p* = 1.48e-06), A930005H10Rik (1.66-fold, *p* = 1.63e-06), 1500015A07Rik (1.61-fold, *p* = 2.09e-06), Xlr3b (1.57-fold, *p* = 2.26e-05), and Me3 (1.57-fold, *p* = 8.47e-07). These genes are involved in metabolism (Adi1, Stbd1, Me3), cilia and flagella functions (Cfap54), and spine morphogenesis and synapse assembly (Xlr3b), with several genes functionally uncharacterized.

**Fig. 5.**
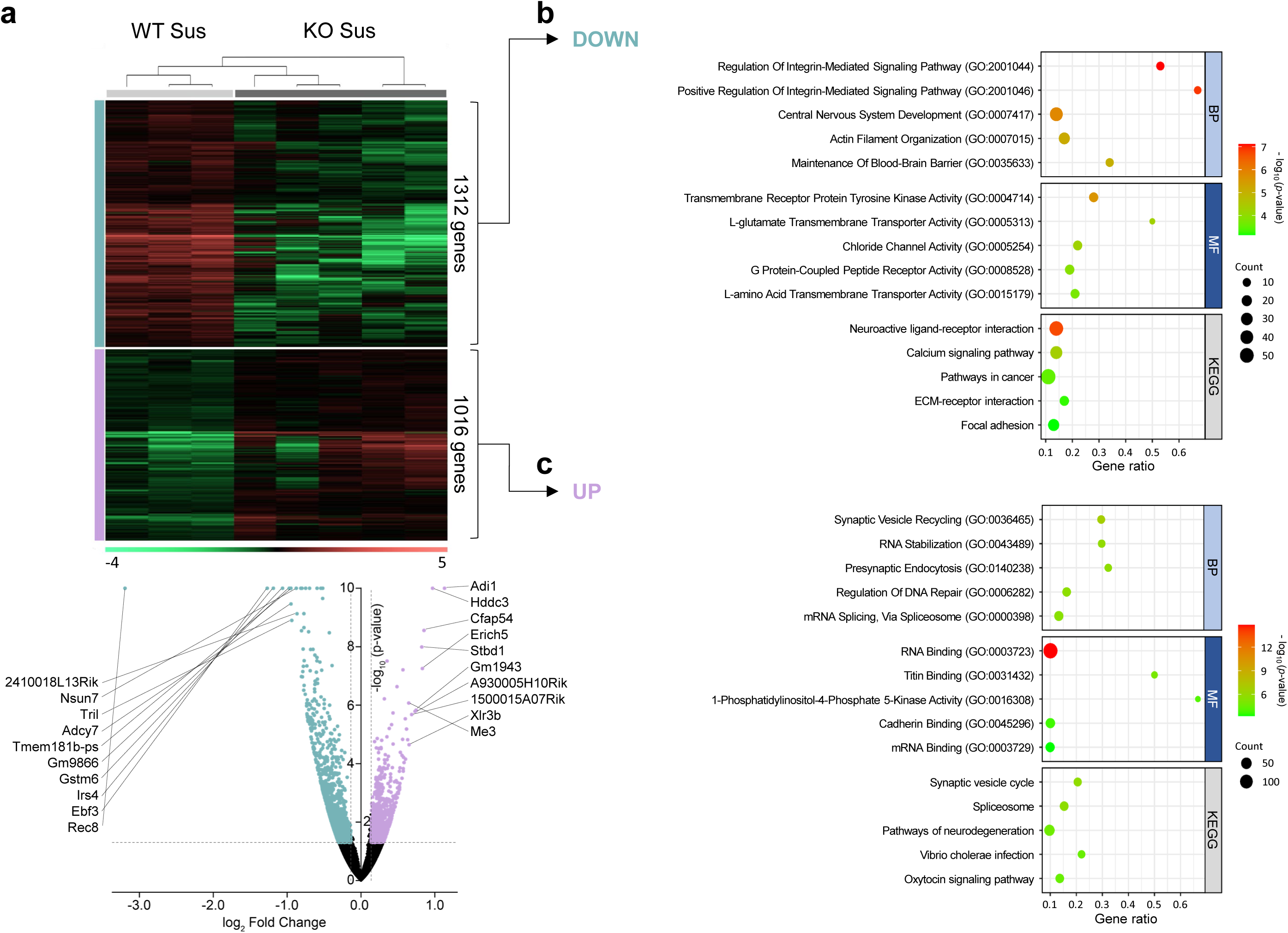
Analysis of SIRT1-KO mice-specific genes in D1-MSNs of susceptible mice. **a** The gene expression differences between wild-type susceptible (WT Sus) and SIRT1-KO susceptible (KO Sus) mice are presented as a heatmap and volcano plot, showing 2328 differentially expressed genes. In the comparison between WT Sus and KO Sus, 1312 genes were decreased and 1016 genes were increased in KO Sus, using thresholds of a fold change of ±1.1 and a *p*-value < 0.05. **b, c** Gene ontology analysis of the downregulated (**b**) and upregulated (**c**) genes was performed using EnrichR, focusing on biological process (BP), KEGG pathway, and molecular function (MF)

The top downregulated genes in KO Sus, ranked by fold change and p-value, were Rec8 (−9.16-fold, *p* ≈ 0), Ebf3 (−2.41-fold, *p* = 1.53e-18), Irs4 (−2.28-fold, *p* = 1.65e-14), Gstm6 (−2.09-fold, *p* = 4.74e-13), Gm9866 (−1.96-fold, *p* = 7.78e-22), Tmem181b-ps (−1.93-fold, *p* = 5.56e-15), Adcy7 (−1.93-fold, *p* = 3.44e-10), Tril (−1.92-fold, *p* = 1.26e-09), Nsun7 (−1.84-fold, *p* = 7.60e-16), and 2410018L13Rik (−1.82-fold, *p* = 7.43e-10). The genes are associated with chromatin remodeling and transcriptional regulation (Rec8, Ebf3), metabolic functions (Gstm6, Nsun7), signal transduction (Irs4, Adcy7), and immune responses (Tril).

The top two enriched terms that were downregulated in the Biological Process GO were Integrin-signaling related terms: ‘Regulation Of Integrin-Mediated Signaling Pathway’ (GO:2001044; −log_10_(*p*) = 7.12) and ‘Positive Regulation Of Integrin-Mediated Signaling Pathway’ (GO:2001046; −log_10_(*p*) = 6.88), both with over 50% gene ratio (Fig. 5b). Other significant downregulated terms included ‘Central Nervous System Development’ (GO:0007417; −log_10_(*p*) = 5.83) and ‘Maintenance Of Blood-Brain Barrier’ (GO:0035633; −log_10_(*p*) = 5.04), indicating potential impacts on development and structural integrity of nervous system. In the Molecular Function GO terms, receptor and channel related terms were detected with the downregulated gene set, such as Transmembrane Receptor Protein Tyrosine Kinase Activity (GO:0004714; −log_10_(*p*) = 5.59), Chloride Channel Activity (GO:0005254; −log_10_(*p*) = 4.26), and G Protein-Coupled Peptide Receptor Activity (GO:0008528; −log_10_(*p*) = 3.91), suggesting potential disruptions in signaling pathways and ion transport mechanisms (Fig. 5b). This is further supported by KEGG analysis, which identified downregulation in ‘Neuroactive ligand-receptor interaction’ (−log_10_(*p*) = 6.73) and ‘Calcium signaling pathway’ (−log_10_(*p*) = 4.35), consistent with the observed disruptions.

On the other hand, several upregulated terms emphasize the changes of synaptic vesicle function and RNA regulation in SIRT1^D1-KO^ susceptible mice (Fig. 5c). Key biological processes and molecular functions include ‘Synaptic Vesicle Recycling’ (GO:0036465; −log_10_(*p*) = 6.78), ‘Presynaptic Endocytosis’ (GO:0140238; −log_10_(*p*) = 5.74), ‘RNA Stabilization’ (GO:0043489; −log_10_(*p*) = 5.86), ‘mRNA Splicing via the Spliceosome’ (GO:0000398; −log_10_(*p*) = 5.49), and ‘RNA Binding’ (GO:0003723; −log_10_(*p*) = 14.8). These findings align with KEGG pathway analyses showing upregulation in the ‘Synaptic Vesicle Cycle’ (−log_10_(*p*) = 5.81), ‘Spliceosome’ (−log_10_(*p*) = 5.69), and ‘Oxytocin Signaling Pathway’ (−log10(*p*) = 4.45), highlighting significant impacts on synaptic function and RNA processing.

## Discussion

Chronic stress can negatively impact brain function, and epigenetic mechanisms like histone modifications are thought to play a key role (Krishnan and Nestler 2008). SIRT1, a histone deacetylase, is a crucial regulator of neuronal plasticity; however, how SIRT1 specifically affects the transcriptional response to stress remains unclear. In previous studies, we analyzed and reported on aspects of SIRT1 function in D1 neurons, primarily focusing on behaviors and neuronal activities related to receptor and synaptic functions (Kim et al. 2024; Kim et al. 2016). While these studies were comprehensive, we did not extend to broader gene expression profiles influenced by Sirt1-KO, nor did they include bioinformatic analyses of stress-susceptible SIRT1-KO mice. To address these gaps, in this study we have refined our RNA sequencing analysis by employing more precise sample selections and threshold adjustments to investigate the broader impacts of CSDS on gene expression in D1 medium spiny neurons of wild-type and SIRT1^D1-KO^ mice, aiming to identify SIRT1-dependent and -independent mechanisms of stress-induced neuronal adaptations. Specifically, we focus on changes in mitochondrial function, circadian rhythm regulation, and histone acetylation to understand how SIRT1 deletion affects stress-induced gene expression changes.

The gene set enrichment analysis (GSEA) revealed significant alterations in pathways related to cellular metabolism and mitochondrial function in D1-MSNs of WT mice following CSDS. After CSDS, we observed significant downregulation in gene sets associated with mitochondrial pathways, such as ‘Cellular Respiration’, ‘Mitochondrial ATP Synthesis Coupled Electron Transport’, and ‘Oxidative Phosphorylation’, in D1-MSNs of wild-type WT susceptible mice. The dysregulation of these pathways suggests that CSDS might lead to a diminished capacity for ATP production, potentially contributing to an energy deficit state within the brain, which is implicated in the development of stress-related disorders (Rezin et al. 2009; Picard, Juster, and McEwen 2014; Filiou and Sandi 2019; Morava and Kozicz 2013). Furthermore, our results indicated alterations in gene sets associated with ‘Energy Derivation By Oxidation Of Organic Compounds’ and ‘Thermogenesis’, supporting the notion that the stress response involves a systemic alteration of metabolic processes (Bartness et al. 2010). Additionally, there was significant enrichment of downregulated genes in datasets related to ‘Active Monoatomic Ion Transmembrane Transporter Activity’ and ‘NADH Dehydrogenase (Ubiquinone) Activity’. This points towards a potential compromise in the cells’ ability to maintain ion gradients and redox balance, which are essential for cellular signaling and synaptic plasticity (Manoli et al. 2007). Collectively, these findings suggest that CSDS leads to a widespread downregulation of genes crucial for mitochondrial function and energy metabolism in D1-MSNs of susceptible WT mice. This is in line with a growing body of research indicating that chronic stress can result in mitochondrial abnormalities, potentially contributing to the pathophysiology of stress-related disorders (Picard, Juster, and McEwen 2014; Rezin et al. 2009).

The GO analysis of DEGs between SIRT1^D1-KO^ control (KO Con) and SIRT1^D1-KO^ susceptible (KO Sus) mice revealed significant enrichment of terms related to circadian rhythm and transcriptional regulation. A significant finding in the KO Con vs. KO Sus results was the increased enrichment of ‘E-box Binding’, which implies a heightened transcriptional drive of circadian genes, likely as a compensatory response to the absence of SIRT1’s regulatory influence as a deacetylase (Hirayama et al. 2007; Nakahata et al. 2008). E-box motifs are essential elements within the promoter regions of circadian clock-controlled genes, recognized and driven by CLOCK:BMAL1 heterodimers. These transcription factors initiate the transcription of negative arm core clock genes, such as Period (PER) and Cryptochromes (CRY), which are crucial for maintaining circadian oscillations (Gekakis et al. 1998; Kondratov et al. 2006). The increased E-box binding activity observed in SIRT1^D1-KO^ mice might reflect an attempt to maintain circadian gene expression in the absence of SIRT1’s modulating effects on CLOCK:BMAL1 activity (Asher et al. 2008). Given that SIRT1 interacts with the circadian core clock machinery by deacetylating key transcription factors like CLOCK (Hirayama et al. 2007), the increased E-box binding activity in SIRT1^D1-KO^ Sus mice suggests a possible compensatory response to maintain circadian gene expression in the absence of SIRT1’s regulation. The heightened binding at E-box motifs in susceptible animals may be indicative of altered transcriptional activity of genes that are critical for the adaptation to environmental stressors. This finding is further supported by the enrichment of ‘Entrainment of Circadian Clock by Photoperiod’ and the ‘Circadian Rhythm’ pathway in KEGG analysis, indicating a systemic reorganization or amplification of the circadian machinery in response to stress and the lack of SIRT1. These findings suggest that SIRT1 plays a crucial role in modulating the circadian clock and photoperiodic responses in the context of stress susceptibility. The circadian clock is a critical regulator of physiological and behavioral processes, and its dysregulation has been implicated in various psychiatric disorders, including depression and anxiety (McClung 2013; Landgraf, McCarthy, and Welsh 2014). The KEGG pathway analysis further supported these findings, revealing that the most significantly enriched term was “Circadian Rhythm”. This suggests that stress and SIRT1 deletion may alter behaviors via the regulation of clock-controlled genes in D1-MSNs. The circadian system is highly interconnected with the stress response, and disruptions in circadian rhythms have been associated with increased vulnerability to stress-related disorders (Logan and McClung 2019; Koch et al. 2017). The dysregulation of circadian genes in SIRT1^D1-KO^ Sus mice may contribute to their heightened susceptibility to chronic stress.

The downregulation of ‘Nucleosome Binding’ and ‘Histone Reader Activity’ in D1-MSNs of SIRT1^D1-KO^ Sus mice suggests that the absence of SIRT1 may lead to altered chromatin accessibility and transcriptional regulation, potentially contributing to maladaptive neuronal responses to CSDS. The decreased ‘Histone Reader Activity’ in SIRT1^D1-KO^ Sus mice further supports the notion that SIRT1 is critical for maintaining proper chromatin organization and transcriptional control. Histone readers are proteins that recognize and bind to specific histone modifications, such as acetylation, methylation, and phosphorylation, and recruit other chromatin-modifying enzymes and transcription factors (Yun et al. 2011). The downregulation of histone reader activity in the absence of SIRT1 may lead to impaired interpretation of histone marks and disrupted recruitment of chromatin regulators, resulting in altered gene expression patterns in response to CSDS. SIRT1’s function as an HDAC is particularly relevant in the context of stress-induced neuronal plasticity and adaptation. SIRT1 deacetylates specific histone residues, such as H3K9 and H4K16, which are associated with transcriptional repression and chromatin condensation (Vaquero et al. 2004; Vaquero et al. 2007). In the absence of SIRT1, increased histone acetylation may lead to an open chromatin state and dysregulated gene expressions involved in stress responses and neuronal function.

In the same regard, significant alterations in histone acetylation are also evident in D1-MSNs of SIRT1^D1-KO^ mice compared to wild-type controls. In D1-MSNs of SIRT1^D1-KO^ mice (WT Con vs. KO Con), the absence of SIRT1 led to a notable increase in gene ontologies associated with histone acetylation activities, specifically, an increase in ‘Histone H4K5 Acetyltransferase Activity’ and ‘Histone H4K8 Acetyltransferase Activity’. The specific enrichment of H4K5 and H4K8 acetyltransferase activities in SIRT1 KO D1-MSNs suggests that SIRT1 may preferentially target these lysine residues for deacetylation. Previous studies have shown that SIRT1 can indeed deacetylate histone H4 at multiple lysine residues, including K5, K8, K12, and K16 (Imai et al. 2000; Vaquero et al. 2004). The deacetylation of H4K5 and H4K8 by SIRT1 has been implicated in the formation of facultative heterochromatin and the repression of gene expression (Vaquero et al. 2004). Therefore, the absence of SIRT1 in D1-MSNs may lead to increased acetylation levels at H4K5 and H4K8, resulting in a more open chromatin structure and potentially increased expression of genes associated with these histone modifications. These findings corroborate the knockout of the SIRT1 gene and may serve as a validation of its functional absence, which is consistent with the known role of SIRT1 in regulating histone acetylation. The upregulation of H4K5 and H4K8 acetyltransferase activities in SIRT1^D1-KO^ D1-MSNs may also have implications for the regulation of specific genes and cellular processes in these neurons. For example, H4K5 acetylation is associated with the activation of genes involved in learning and memory, such as BDNF and FOS (Peleg et al. 2010). Additionally, H4K8 acetylation has been implicated in the regulation of cell cycle progression and cellular differentiation (Sobel et al. 1995). Thus, the increased expression of genes with H4K5 and H4K8 acetyltransferase activities in SIRT1^D1-KO^ D1-MSNs may lead to alterations in the expression of specific genes and cellular processes related to these histone modifications.

In the comparison between wild-type Sus and SIRT1^D1-KO^ Sus mice, the downregulated DEGs significantly matched with integrin, receptor tyrosine kinase, and G-protein coupled receptor (GPCR) signaling pathways, indicating potential impairments in cell adhesion and communication in SIRT1^D1-KO^ Sus mice. Downregulated terms also included ‘Central Nervous System Development’ and ‘Maintenance Of Blood-Brain Barrier’, suggesting adverse effects on neural development and barrier integrity. It is noteworthy that blood-brain barrier integrity is closely associated with depression (Menard et al. 2017; Dudek et al. 2020).

Conversely, upregulated terms in SIRT1^D1-KO^ Sus mice highlighted enhanced synaptic activity and RNA processing, with significant terms such as ‘Synaptic Vesicle Recycling’, ‘Presynaptic Endocytosis’, and ‘RNA Stabilization’. KEGG pathway analysis confirmed upregulation in the ‘Synaptic Vesicle Cycle’ and ‘Spliceosome,’ indicating increased synaptic vesicle turnover and RNA splicing. These changes suggest compensatory mechanisms aimed at maintaining synaptic function and gene regulation in the absence of SIRT1. Additionally, KEGG analysis showed decreased ‘Neuroactive ligand-receptor interaction’ and ‘Calcium signaling pathway’ activities, potentially compromising neurotransmission and cellular communication.

Interestingly, many of the top-regulated genes overlapped in both KO Con (vs WT Con) and KO Sus (vs WT Sus) with the same direction of gene expression changes. This alignment is also seen in overlapping GO terms, such as decreased ‘Neuroactive ligand-receptor interaction’ and other channel and GPCR-mediated signaling, as well as increased synaptic assembly and vesicle cycling. To provide an overview of the differences between groups, DEGs from each comparison were illustrated in a Venn diagram (Fig. 6a). When comparing DEGs between each group, we observed minimal overlap between WT Sus (vs Con) and KO Sus (vs Con) (blue dotted outline in Fig. 6A). This suggests that different pathways are activated in SIRT1^D1-KO^ D1-neurons, leading to susceptibility. Notably, the relatively low number of DEGs in KO Sus implicates SIRT1 as the main player inducing susceptibility in wild-type D1 MSNs. Previous research (Kim et al. 2024) also showed that SIRT1^D1-KO^ mice exhibit more resilient phenotypes compared to their wild-type littermates. Additionally, we found that SIRT1 robustly modulates synaptic genes in WT Sus (vs Con) DEGs; whereas, in this study, SIRT1^D1-KO^ Con and Sus display distinct results in GO analysis compared to the wild-type, including circadian clock pathways and cellular structure, which need further investigation. Although there is a substantial overlap in shared genes between KO Con (vs WT Con) and KO Sus (vs WT Sus), KO Sus exhibits a significant number of unique differentially expressed genes (1589 genes; red dotted encirclement in Fig 6a). Furthermore, KO Con (vs WT Con) and KO Sus (vs WT Sus) show significant overlaps with 310 uniquely shared genes (green dotted encirclement in Fig 6a), potentially modulated by SIRT1. Notably, these 310 shared DEGs expressed in SIRT1^D1-KO^ mice are highly enriched genes for depression-related terms in a human disease-oriented GO domain, DisGeNET (Fig. 6a, inset). This highlights the need for further investigation into the differential regulation of these genes in control and susceptible mice to better understand their roles and SIRT1 actions in depression.

**Fig. 6.**
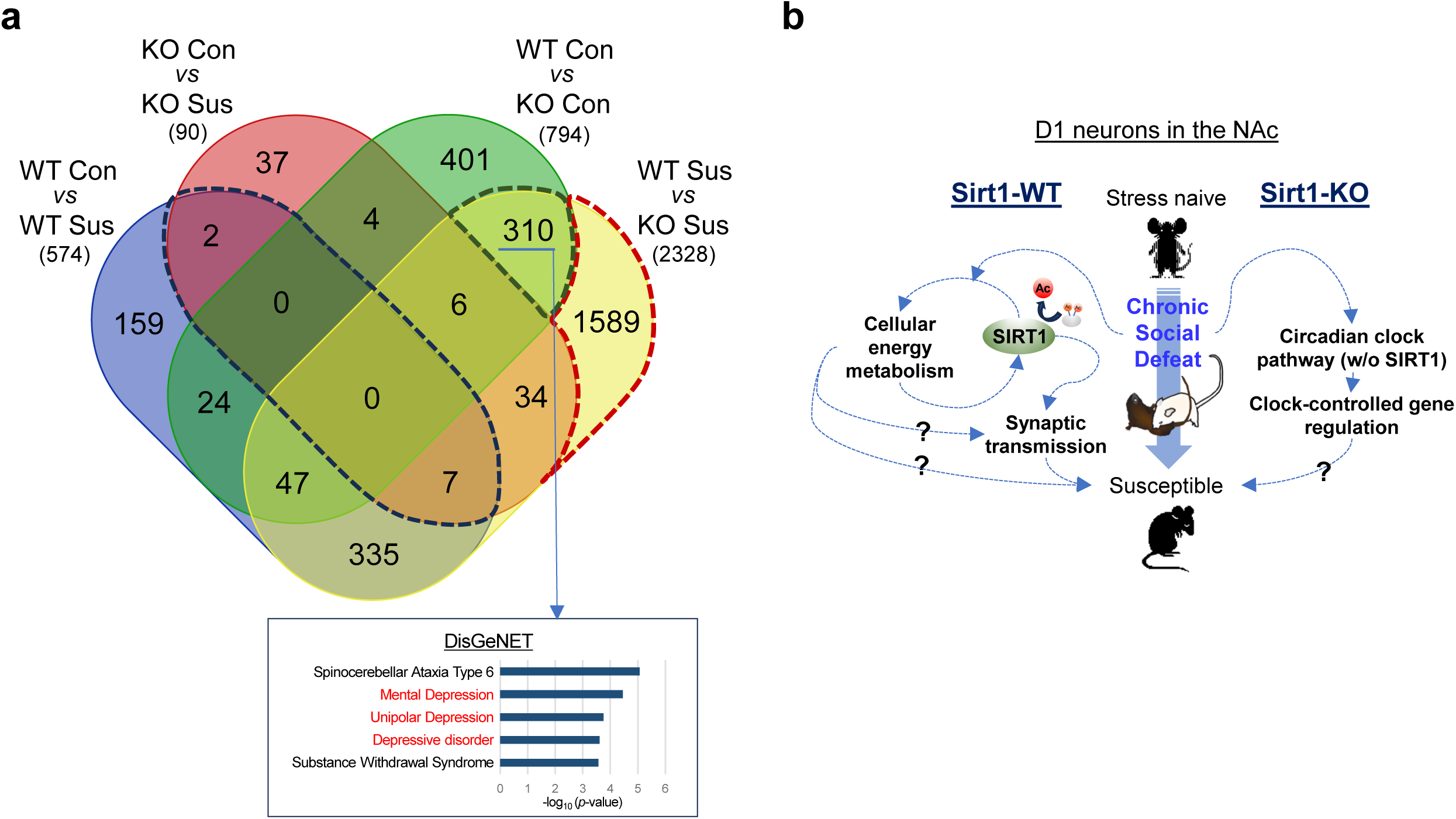
Comparisons of differentially expressed gene sets and proposed model. **a** Comprehensive analysis of all groups (WT Con, WT Sus, KO Con, and KO Sus) was performed using a Venn diagram to identify overarching patterns and differences. The 310 overlapping DEGs from the comparisons of KO Con (vs WT Con) and KO Sus (vs WT Sus) were further analyzed via gene enrichment analysis (DisGeNET; inset). **b** A proposed model of SIRT1-dependent and -independent mechanisms of depression in D1-MSNs. In wild-type mice, SIRT1 and related cellular metabolism pathways contribute to susceptibility. Conversely, in SIRT1^D1-KO^ mice, the stress susceptibility may be induced by circadian clock-mediated transcriptional regulation in D1-MSNs

In summary, the genes related to cellular energy metabolism and deacetylase function of SIRT1 are important in eliciting behavioral deficits in wild-type mice. On the other hand, the absence of functional SIRT1 evokes alternative pathways, such as circadian clock machinery or other signaling pathways, upon chronic stress and induces similar depression like phenotypes in SIRT1^D1-KO^ mice (Fig. 6b). Further studies investigating the particular genes and signaling pathways regulated by SIRT1-mediated histone deacetylation in D1-MSNs may provide valuable insights into the molecular underpinnings of depression and mood disorders. Overall, these findings suggest that different molecular pathways can induce behavioral susceptibility, highlighting the heterogeneous pathophysiology of depression, and each pathway may represent a promising therapeutic target for the condition.

## Statements and Declarations

### Author Contributions

H-D.K., H.J.C., S.Q., and D.F. designed the research, analyzed data, and wrote the manuscript. H-D.K., A.N., and A.B. performed behavioral tests. H-D.K., H.J.C., and D.F. contributed bioinformatic analysis. H-D.K. performed all other experiments.

### Ethics Approval

All animal procedures were approved by the University of Arizona Institutional Animal Care and Use Committees.

### Data Availability

RNA-seq data have been deposited to the NCBI Gene Expression Omnibus under accession number GSE147640.

## References

Abe-Higuchi, N., S. Uchida, H. Yamagata, F. Higuchi, T. Hobara, K. Hara, A. Kobayashi, and Y. Watanabe. 2016. ‘Hippocampal Sirtuin 1 Signaling Mediates Depression-like Behavior’, Biol Psychiatry, 80: 815–26.

Asher, G., D. Gatfield, M. Stratmann, H. Reinke, C. Dibner, F. Kreppel, R. Mostoslavsky, F. W. Alt, and U. Schibler. 2008. ‘SIRT1 regulates circadian clock gene expression through PER2 deacetylation’, Cell, 134: 317–28.

Bartness, T. J., Y. B. Shrestha, C. H. Vaughan, G. J. Schwartz, and C. K. Song. 2010. ‘Sensory and sympathetic nervous system control of white adipose tissue lipolysis’, Mol Cell Endocrinol, 318: 34–43.

Berridge, M. J. 2012. ‘Calcium signalling remodelling and disease’, Biochem Soc Trans, 40: 297–309.

Breuer, M. E., W. J. Koopman, S. Koene, M. Nooteboom, R. J. Rodenburg, P. H. Willems, and J. A. Smeitink. 2013. ‘The role of mitochondrial OXPHOS dysfunction in the development of neurologic diseases’, Neurobiol Dis, 51: 27–34.

Catterall, W. A. 2011. ‘Voltage-gated calcium channels’, Cold Spring Harb Perspect Biol, 3: a003947.

consortium, Converge. 2015. ‘Sparse whole-genome sequencing identifies two loci for major depressive disorder’, Nature, 523: 588–91.

Dudek, K. A., L. Dion-Albert, M. Lebel, K. LeClair, S. Labrecque, E. Tuck, C. Ferrer Perez, S. A. Golden, C. Tamminga, G. Turecki, N. Mechawar, S. J. Russo, and C. Menard. 2020. ‘Molecular adaptations of the blood-brain barrier promote stress resilience vs. depression’, Proc Natl Acad Sci U S A, 117: 3326–36.

Fava, M., and K. G. Davidson. 1996. ‘Definition and epidemiology of treatment-resistant depression’, Psychiatr Clin North Am, 19: 179–200.

Ferguson, D., J. W. Koo, J. Feng, E. Heller, J. Rabkin, M. Heshmati, W. Renthal, R. Neve, X. Liu, N. Shao, V. Sartorelli, L. Shen, and E. J. Nestler. 2013. ‘Essential role of SIRT1 signaling in the nucleus accumbens in cocaine and morphine action’, J Neurosci, 33: 16088–98.

Fields, R. D. 2008. ‘White matter in learning, cognition and psychiatric disorders’, Trends Neurosci, 31: 361–70.

Filiou, M. D., and C. Sandi. 2019. ‘Anxiety and Brain Mitochondria: A Bidirectional Crosstalk’, Trends Neurosci, 42: 573–88.

Garner, C. C., R. G. Zhai, E. D. Gundelfinger, and N. E. Ziv. 2002. ‘Molecular mechanisms of CNS synaptogenesis’, Trends Neurosci, 25: 243–51.

Gekakis, N., D. Staknis, H. B. Nguyen, F. C. Davis, L. D. Wilsbacher, D. P. King, J. S. Takahashi, and C. J. Weitz. 1998. ‘Role of the CLOCK protein in the mammalian circadian mechanism’, Science, 280: 1564–9.

Golden, S. A., H. E. Covington, 3rd, O. Berton, and S. J. Russo. 2011. ‘A standardized protocol for repeated social defeat stress in mice’, Nat Protoc, 6: 1183–91.

Gong, S., M. Doughty, C. R. Harbaugh, A. Cummins, M. E. Hatten, N. Heintz, and C. R. Gerfen. 2007. ‘Targeting Cre recombinase to specific neuron populations with bacterial artificial chromosome constructs’, J Neurosci, 27: 9817–23.

Herskovits, A. Z., and L. Guarente. 2014. ‘SIRT1 in neurodevelopment and brain senescence’, Neuron, 81: 471–83.

Hirayama, J., S. Sahar, B. Grimaldi, T. Tamaru, K. Takamatsu, Y. Nakahata, and P. Sassone-Corsi. 2007. ‘CLOCK-mediated acetylation of BMAL1 controls circadian function’, Nature, 450: 1086–90.

Hisahara, S., S. Chiba, H. Matsumoto, M. Tanno, H. Yagi, S. Shimohama, M. Sato, and Y. Horio. 2008. ‘Histone deacetylase SIRT1 modulates neuronal differentiation by its nuclear translocation’, Proc Natl Acad Sci U S A, 105: 15599–604.

Imai, S., C. M. Armstrong, M. Kaeberlein, and L. Guarente. 2000. ‘Transcriptional silencing and longevity protein Sir2 is an NAD-dependent histone deacetylase’, Nature, 403: 795–800.

Jenuwein, T., and C. D. Allis. 2001. ‘Translating the histone code’, Science, 293: 1074–80.

Kessler, R. C., M. Angermeyer, J. C. Anthony, D. E. Graaf R, K. Demyttenaere, I. Gasquet, D. E. Girolamo G, S. Gluzman, O. Gureje, J. M. Haro, N. Kawakami, A. Karam, D. Levinson, M. E. Medina Mora, M. A. Oakley Browne, J. Posada-Villa, D. J. Stein, C. H. Adley Tsang, S. Aguilar-Gaxiola, J. Alonso, S. Lee, S. Heeringa, B. E. Pennell, P. Berglund, M. J. Gruber, M. Petukhova, S. Chatterji, and T. B. Ustun. 2007. ‘Lifetime prevalence and age-of-onset distributions of mental disorders in the World Health Organization’s World Mental Health Survey Initiative’, World Psychiatry, 6: 168–76.

Kim, H. D., J. Hesterman, T. Call, S. Magazu, E. Keeley, K. Armenta, H. Kronman, R. L. Neve, E. J. Nestler, and D. Ferguson. 2016. ‘SIRT1 Mediates Depression-Like Behaviors in the Nucleus Accumbens’, J Neurosci, 36: 8441–52.

Kim, H. D., J. Wei, T. Call, X. Ma, N. T. Quintus, A. J. Summers, S. Carotenuto, R. Johnson, A. Nguyen, Y. Cui, J. G. Park, S. Qiu, and D. Ferguson. 2024. ‘SIRT1 Coordinates Transcriptional Regulation of Neural Activity and Modulates Depression-Like Behaviors in the Nucleus Accumbens’, Biol Psychiatry.

Kim, H. D., J. Wei, T. Call, N. T. Quintus, A. J. Summers, S. Carotenuto, R. Johnson, X. Ma, C. Xu, J. G. Park, S. Qiu, and D. Ferguson. 2021. ‘Shisa6 mediates cell-type specific regulation of depression in the nucleus accumbens’, Mol Psychiatry, 26: 7316–27.

Kim, Hee-Dae, Tanessa Call, Samantha Carotenuto, Ross Johnson, and Deveroux Ferguson. 2017. ‘Testing Depression in Mice: a Chronic Social Defeat Stress Model’, Bio Protoc, 7.

Koch, C. E., B. Leinweber, B. C. Drengberg, C. Blaum, and H. Oster. 2017. ‘Interaction between circadian rhythms and stress’, Neurobiol Stress, 6: 57–67.

Kondratov, R. V., A. A. Kondratova, V. Y. Gorbacheva, O. V. Vykhovanets, and M. P. Antoch. 2006. ‘Early aging and age-related pathologies in mice deficient in BMAL1, the core componentof the circadian clock’, Genes Dev, 20: 1868–73.

Koo, J. W., B. Labonte, O. Engmann, E. S. Calipari, B. Juarez, Z. Lorsch, J. J. Walsh, A. K. Friedman, J. T. Yorgason, M. H. Han, and E. J. Nestler. 2016. ‘Essential Role of Mesolimbic Brain-Derived Neurotrophic Factor in Chronic Social Stress-Induced Depressive Behaviors’, Biol Psychiatry, 80: 469–78.

Krishnan, V., M. H. Han, D. L. Graham, O. Berton, W. Renthal, S. J. Russo, Q. Laplant, A. Graham, M. Lutter, D. C. Lagace, S. Ghose, R. Reister, P. Tannous, T. A. Green, R. L. Neve, S. Chakravarty, A. Kumar, A. J. Eisch, D. W. Self, F. S. Lee, C. A. Tamminga, D. C. Cooper, H. K. Gershenfeld, and E. J. Nestler. 2007. ‘Molecular adaptations underlying susceptibility and resistance to social defeat in brain reward regions’, Cell, 131: 391–404.

Krishnan, V., and E. J. Nestler. 2008. ‘The molecular neurobiology of depression’, Nature, 455: 894–902.

Kuleshov, M. V., M. R. Jones, A. D. Rouillard, N. F. Fernandez, Q. Duan, Z. Wang, S. Koplev, S. L. Jenkins, K. M. Jagodnik, A. Lachmann, M. G. McDermott, C. D. Monteiro, G. W. Gundersen, and A. Ma’ayan. 2016. ‘Enrichr: a comprehensive gene set enrichment analysis web server 2016 update’, Nucleic Acids Res, 44: W90–7.

Landgraf, D., J. E. Long, C. D. Proulx, R. Barandas, R. Malinow, and D. K. Welsh. 2016. ‘Genetic Disruption of Circadian Rhythms in the Suprachiasmatic Nucleus Causes Helplessness, Behavioral Despair, and Anxiety-like Behavior in Mice’, Biol Psychiatry, 80: 827–35.

Landgraf, D., M. J. McCarthy, and D. K. Welsh. 2014. ‘Circadian clock and stress interactions in the molecular biology of psychiatric disorders’, Curr Psychiatry Rep, 16: 483.

Li, H., G. K. Rajendran, N. Liu, C. Ware, B. P. Rubin, and Y. Gu. 2007. ‘SirT1 modulates the estrogen-insulin-like growth factor-1 signaling for postnatal development of mammary gland in mice’, Breast Cancer Res, 9: R1.

Li, J. Z., B. G. Bunney, F. Meng, M. H. Hagenauer, D. M. Walsh, M. P. Vawter, S. J. Evans, P. V. Choudary, P. Cartagena, J. D. Barchas, A. F. Schatzberg, E. G. Jones, R. M. Myers, S. J. Watson, Jr., H. Akil, and W. E. Bunney. 2013. ‘Circadian patterns of gene expression in the human brain and disruption in major depressive disorder’, Proc Natl Acad Sci U S A, 110: 9950–5.

Libert, S., K. Pointer, E. L. Bell, A. Das, D. E. Cohen, J. M. Asara, K. Kapur, S. Bergmann, M. Preisig, T. Otowa, K. S. Kendler, X. Chen, J. M. Hettema, E. J. van den Oord, J. P. Rubio, and L. Guarente. 2011. ‘SIRT1 activates MAO-A in the brain to mediate anxiety and exploratory drive’, Cell, 147: 1459–72.

Logan, R. W., and C. A. McClung. 2019. ‘Rhythms of life: circadian disruption and brain disorders across the lifespan’, Nat Rev Neurosci, 20: 49–65.

Lu, G., J. Li, H. Zhang, X. Zhao, L. J. Yan, and X. Yang. 2018. ‘Role and Possible Mechanisms of Sirt1 in Depression’, Oxid Med Cell Longev, 2018: 8596903.

Manoli, I., S. Alesci, M. R. Blackman, Y. A. Su, O. M. Rennert, and G. P. Chrousos. 2007. ‘Mitochondria as key components of the stress response’, Trends Endocrinol Metab, 18: 190–8.

McCarthy, M. J., H. Wei, C. M. Nievergelt, A. Stautland, A. X. Maihofer, D. K. Welsh, P. Shilling, M. Alda, N. Alliey-Rodriguez, A. Anand, O. A. Andreasson, Y. Balaraman, W. H. Berrettini, H. Bertram, K. J. Brennand, J. R. Calabrese, C. V. Calkin, A. Claasen, C. Conroy, W. H. Coryell, D. W. Craig, N. D’Arcangelo, A. Demodena, S. Djurovic, S. Feeder, C. Fisher, N. Frazier, M. A. Frye, F. H. Gage, K. Gao, J. Garnham, E. S. Gershon, K. Glazer, F. Goes, T. Goto, G. Harrington, P. Jakobsen, M. Kamali, E. Karberg, M. Kelly, S. G. Leckband, F. Lohoff, M. G. McInnis, F. Mondimore, G. Morken, J. I. Nurnberger, S. Obral, K. J. Oedegaard, A. Ortiz, M. Ritchey, K. Ryan, M. Schinagle, H. Schoeyen, C. Schwebel, M. Shaw, T. Shekhtman, C. Slaney, E. Stapp, S. Szelinger, B. Tarwater, P. P. Zandi, and J. R. Kelsoe. 2019. ‘Chronotype and cellular circadian rhythms predict the clinical response to lithium maintenance treatment in patients with bipolar disorder’, Neuropsychopharmacology, 44: 620–28.

McClung, C. A. 2013. ‘How might circadian rhythms control mood? Let me count the ways’, Biol Psychiatry, 74: 242–9.

Menard, C., M. L. Pfau, G. E. Hodes, V. Kana, V. X. Wang, S. Bouchard, A. Takahashi, M. E. Flanigan, H. Aleyasin, K. B. LeClair, W. G. Janssen, B. Labonte, E. M. Parise, Z. S. Lorsch, S. A. Golden, M. Heshmati, C. Tamminga, G. Turecki, M. Campbell, Z. A. Fayad, C. Y. Tang, M. Merad, and S. J. Russo. 2017. ‘Social stress induces neurovascular pathology promoting depression’, Nat Neurosci, 20: 1752–60.

Michan, S., Y. Li, M. M. Chou, E. Parrella, H. Ge, J. M. Long, J. S. Allard, K. Lewis, M. Miller, W. Xu, R. F. Mervis, J. Chen, K. I. Guerin, L. E. Smith, M. W. McBurney, D. A. Sinclair, M. Baudry, R. de Cabo, and V. D. Longo. 2010. ‘SIRT1 is essential for normal cognitive function and synaptic plasticity’, J Neurosci, 30: 9695–707.

Morava, E., and T. Kozicz. 2013. ‘Mitochondria and the economy of stress (mal)adaptation’, Neurosci Biobehav Rev, 37: 668–80.

Nakahata, Y., M. Kaluzova, B. Grimaldi, S. Sahar, J. Hirayama, D. Chen, L. P. Guarente, and P. Sassone-Corsi. 2008. ‘The NAD+-dependent deacetylase SIRT1 modulates CLOCK-mediated chromatin remodeling and circadian control’, Cell, 134: 329–40.

Oberdoerffer, P., S. Michan, M. McVay, R. Mostoslavsky, J. Vann, S. K. Park, A. Hartlerode, J. Stegmuller, A. Hafner, P. Loerch, S. M. Wright, K. D. Mills, A. Bonni, B. A. Yankner, R. Scully, T. A. Prolla, F. W. Alt, and D. A. Sinclair. 2008. ‘SIRT1 redistribution on chromatin promotes genomic stability but alters gene expression during aging’, Cell, 135: 907–18.

Peleg, S., F. Sananbenesi, A. Zovoilis, S. Burkhardt, S. Bahari-Javan, R. C. Agis-Balboa, P. Cota, J. L. Wittnam, A. Gogol-Doering, L. Opitz, G. Salinas-Riester, M. Dettenhofer, H. Kang, L. Farinelli, W. Chen, and A. Fischer. 2010. ‘Altered histone acetylation is associated with age-dependent memory impairment in mice’, Science, 328: 753–6.

Picard, M., R. P. Juster, and B. S. McEwen. 2014. ‘Mitochondrial allostatic load puts the ‘gluc’ back in glucocorticoids’, Nat Rev Endocrinol, 10: 303–10.

Renthal, W., A. Kumar, G. Xiao, M. Wilkinson, H. E. Covington, 3rd, I. Maze, D. Sikder, A. J. Robison, Q. LaPlant, D. M. Dietz, S. J. Russo, V. Vialou, S. Chakravarty, T. J. Kodadek, A. Stack, M. Kabbaj, and E. J. Nestler. 2009. ‘Genome-wide analysis of chromatin regulation by cocaine reveals a role for sirtuins’, Neuron, 62: 335–48.

Rezin, G. T., G. Amboni, A. I. Zugno, J. Quevedo, and E. L. Streck. 2009. ‘Mitochondrial dysfunction and psychiatric disorders’, Neurochem Res, 34: 1021–9.

Rush, A. J., M. H. Trivedi, S. R. Wisniewski, A. A. Nierenberg, J. W. Stewart, D. Warden, G. Niederehe, M. E. Thase, P. W. Lavori, B. D. Lebowitz, P. J. McGrath, J. F. Rosenbaum, H. A. Sackeim, D. J. Kupfer, J. Luther, and M. Fava. 2006. ‘Acute and longer-term outcomes in depressed outpatients requiring one or several treatment steps: a STAR*D report’, Am J Psychiatry, 163: 1905–17.

Sanz, E., L. Yang, T. Su, D. R. Morris, G. S. McKnight, and P. S. Amieux. 2009. ‘Cell-type-specific isolation of ribosome-associated mRNA from complex tissues’, Proc Natl Acad Sci U S A, 106: 13939–44.

Schwacha, A., and N. Kleckner. 1995. ‘Identification of double Holliday junctions as intermediates in meiotic recombination’, Cell, 83: 783–91.

Shapiro, L., J. Love, and D. R. Colman. 2007. ‘Adhesion molecules in the nervous system: structural insights into function and diversity’, Annu Rev Neurosci, 30: 451–74.

Sobel, R. E., R. G. Cook, C. A. Perry, A. T. Annunziato, and C. D. Allis. 1995. ‘Conservation of deposition-related acetylation sites in newly synthesized histones H3 and H4’, Proc Natl Acad Sci U S A, 92: 1237–41.

Tang, D., M. Chen, X. Huang, G. Zhang, L. Zeng, G. Zhang, S. Wu, and Y. Wang. 2023. ‘SRplot: A free online platform for data visualization and graphing’, PLoS One, 18: e0294236.

Tran, T. S., A. L. Kolodkin, and R. Bharadwaj. 2007. ‘Semaphorin regulation of cellular morphology’, Annu Rev Cell Dev Biol, 23: 263–92.

Trivedi, M. H., A. J. Rush, S. R. Wisniewski, A. A. Nierenberg, D. Warden, L. Ritz, G. Norquist, R. H. Howland, B. Lebowitz, P. J. McGrath, K. Shores-Wilson, M. M. Biggs, G. K. Balasubramani, M. Fava, and Star D. Study Team. 2006. ‘Evaluation of outcomes with citalopram for depression using measurement-based care in STAR*D: implications for clinical practice’, Am J Psychiatry, 163: 28–40.

Vaquero, A., M. Scher, H. Erdjument-Bromage, P. Tempst, L. Serrano, and D. Reinberg. 2007. ‘SIRT1 regulates the histone methyl-transferase SUV39H1 during heterochromatin formation’, Nature, 450: 440–4.

Vaquero, A., M. Scher, D. Lee, H. Erdjument-Bromage, P. Tempst, and D. Reinberg. 2004. ‘Human SirT1 interacts with histone H1 and promotes formation of facultative heterochromatin’, Mol Cell, 16: 93–105.

Yun, M., J. Wu, J. L. Workman, and B. Li. 2011. ‘Readers of histone modifications’, Cell Res, 21: 564–78.

